# TOMM40 Knockdown in Macrophages Inhibits Oxidized LDL-induced NLRP3 Activation and Promotes LXR-β Mediated Cholesterol Transporter Gene Expression

**DOI:** 10.1101/2025.07.01.662675

**Authors:** Sai S. Chelluri, Neil V. Yang, Elizabeth Theusch, Ronald M. Krauss

## Abstract

Atherogenesis has been shown in mice to be dependent on activation of the NLRP3 inflammasome, a cytosolic innate immune sensor activated by a broad range of pathogen and damage associated molecular patterns, including oxidized LDL (oxLDL) in atherosclerosis. Previous work from our group has shown that knockdown of Translocase of Outer Mitochondrial Membrane 40 (*TOMM40),* which encodes a component of the mitochondrial importer TOM complex, increases expression and activity of the nuclear receptor-family transcription factor, liver X receptor (LXR) in hepatocytes. As LXR agonists have been shown to inhibit NLRP3 activation, we confirmed the prediction that *TOMM40* knockdown has this effect in THP-1 monocyte-derived macrophages. Further, we demonstrated that *TOMM40* KD upregulates LXR-mediated macrophage expression of the *ABCA1* and *ABCG1* genes which encode transporters that promote cellular cholesterol efflux, the first step in reverse cholesterol transport to the liver. Taken together, these findings identify a novel mechanism whereby increased LXR activity induced by suppression of *TOMM40* expression in macrophages may retard atherogenesis both by inhibiting inflammation and promoting reverse cholesterol transport.

## Introduction

As a disease of chronic inflammation, atherosclerosis is characterized by the recruitment of monocytes to the arterial wall, where they migrate through the endothelium, proliferate, and differentiate into macrophages^1,2^. As these macrophages take up minimally-modified forms of LDL, such as oxidized-LDL (oxLDL), present in greater abundance under atherogenic conditions, they release pro-inflammatory cytokines which further drive the formation of atherosclerotic lesions^2^. While the impact of oxLDL on cholesterol homeostasis in macrophages is well-established, more recent studies have shown that oxLDL also activates the NLRP3 inflammasome, and that NLRP3 activation is necessary for atherogenesis in mice^3,4,5,6^.

A member of the NOD-like receptor (NLR) family of inflammasomes, NLRP3 is a cytosolic immune multiprotein complex activated by a wide range of pathogen and damage-associated molecular patterns including extracellular ATP^7^, uric acid crystals^8^, and cholesterol crystals^3,4,9^. NLRP3 activation can occur in response to many diverse stimuli, and downstream ion effluxes, lysosomal destabilization, and reactive oxygen species production are considered likely causes of activation rather than binding of any individual ligand^9–11^. NLRP3 activation occurs as a two-step process in which an initial priming signal first increases expression of NLRP3 and pro-IL-1β in a nuclear factor-κB (NF-κB)-dependent manner^6,10,12^. A second activating signal, for example lipopolysaccharide (LPS), then causes assembly in which NLRP3 oligomerizes and recruits the adapter protein apoptosis speck-like protein containing a CARD (ASC), which then recruits pro-caspase-1. Upon clustering, pro-caspase-1 autoproteolytically matures into active caspase-1 and processes cytosolic targets including pro-IL-1β and pro-IL-18 for release from the cell^9,13–15^.

The nuclear receptor-family transcription factors, liver X receptor-α and β (LXR-α/LXR-β), repress inflammatory genes and inhibit NF-κB-mediated transcription^16–18^. Concordantly, LXR agonists exert potent anti-inflammatory and anti-atherogenic effect in mouse models^17,19,20^. LXRs are activated by binding of their natural ligands, in particular sterol metabolites such as 22-(*R*)-hydroxycholesterol, 24-(*S*)-hydroxycholesterol, 27-hydroxycholesterol and 24-(*S*),25-epoxycholesterol^18,21–23^. In addition to their effects antagonizing pro-inflammatory gene expression, LXRs play a critical role in whole body cholesterol homeostasis by inducing tissue-specific gene expression patterns^18,24^. In macrophages, LXRs induce expression of the genes encoding ATP-binding cassette transporter A1 (ABCA1) and ATP binding cassette subfamily G member 1 (ABCG1), which are the main mediators of cholesterol efflux from macrophages, the first step of reverse cholesterol transport to the liver, and mice deficient in each have been shown to develop increased and accelerated atherosclerosis^25–30^.

Our group has previously shown that LXR-β is upregulated and LXR activity is increased due to greater intracellular oxysterol concentration following knockdown in hepatocytes of Translocase of Outer Mitochondrial Membrane 40 (*TOMM40)*^31^*. TOMM40* encodes the critical TOM40 subunit of the mitochondrial TOM complex, which serves to import mitochondrial preproteins synthesized by cytosolic ribosomes^32,33^. While the robust LXR upregulation by *TOMM40* knockdown in hepatocytes suggests that it may result in an anti-inflammatory effect, its direct impact on inflammasome activation and cholesterol metabolism in macrophages have not yet been investigated.

In this study, we demonstrate that *TOMM40* knockdown (KD) inhibits NLRP3 activation in response to pro-inflammatory stimuli such as oxLDL by inhibiting upregulation of NLRP3-pathway components. Further, we show that *TOMM40* knockdown in macrophages upregulates cholesterol transporter genes *ABCA1* and *ABCG1* in an LXR-β-dependent manner.

## Methods

### THP-1 Cell Culture & NLRP3 Activation Assay

THP-1 cells were purchased from the ATCC and cultured in complete RPMI (Gibco) medium (10% FBS, 2.25% glucose, 1% Pen-Strep, 1% Sodium Pyruvate, 1% 1M HEPES) supplemented with 0.1% 2-mercaptoethanol every other day. Cells were tested routinely for mycoplasma by PCR kit (ATCC). To test for NLRP3 activation, THP-1 cells were differentiated into macrophages by incubation with 10 ng/ml phorbol-12-myristate-13-acetate (PMA) before reverse transfection with *TOMM40*-targeting or non-targeting control siRNAs (Invitrogen, s20449 and s79125 respectively) using Lipofectamine RNAiMAX transfection reagent (Invitrogen) and Opti-MEM 1 (Gibco). After 44 hours, the cells were primed with 500 ng/ml – 1 ug/ml bacterial lipopolysaccharide (LPS) added directly to the transfection media for 4 hours. Cells were then stimulated with 10 - 40 ug/ml oxLDL (Invitrogen) for 24 hours or 1 - 5uM Nigericin (Invivogen) for 2 hours in fresh medium. If LXRα/β activity was assessed, 10uM GSK2033 (Millipore-Sigma) was added to the transfection media 24 hours after transfection (20 hours before priming). Supernatants were collected and frozen at -80°C, and cells were washed with PBS before being lysed in complete RLT buffer (Qiagen, 1% 2-mercaptoethanol) for RNA extraction, or in CelLytic M buffer (Sigma-Aldrich) supplemented with Halt Protease Inhibitor Cocktail (Thermo-Fisher) for immunoblotting.

### RT-qPCR

Cellular RNA was extracted with the RNeasy Mini QIAcube Kit (Qiagen) using the QIAcube Connect (Qiagen) according to manufacturer’s protocol. cDNA synthesis was performed on total RNA with the High Capacity cDNA Reverse Transcription kit (Applied Biosystems). mRNA transcript levels were quantified by RT-qPCR using primers from Elim Biopharmaceuticals and SYBR™ Green qPCR Master Mix (Thermo Fisher Scientific) on an ABI PRISM 7900 Sequence Detection System. Primers were as follows:

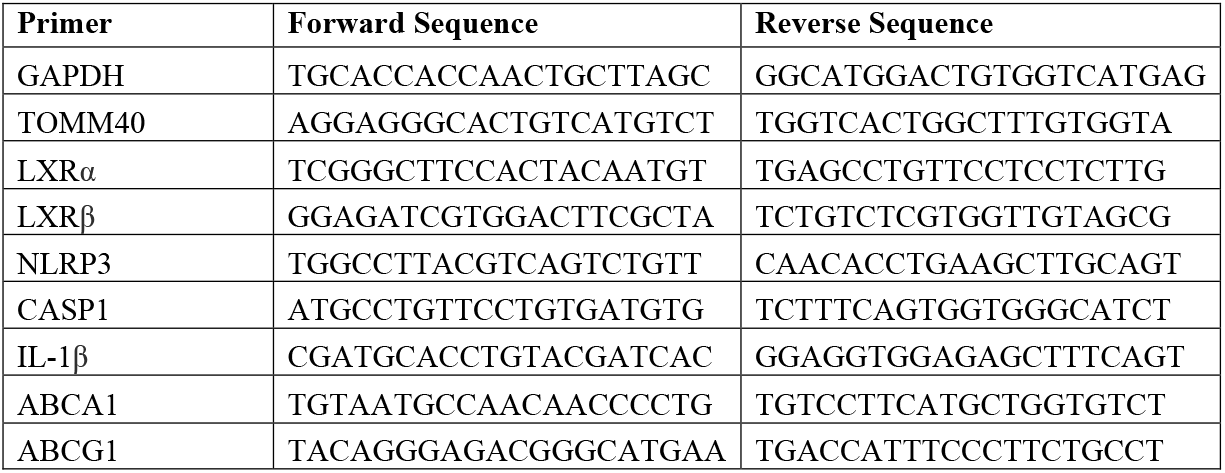

The mean Ct value of triplicates for each sample was normalized to GAPDH as a reference gene. The ΔΔCt method was used to calculate fold changes in transcript levels.

### Cytokine Quantification

Cytokine levels in cell supernatants were assessed using the human IL-1β/IL-1F2 DuoSet ELISA kit (R&D Systems), human IL-18 DuoSet ELISA kit (R&D Systems), and human TNF-α DuoSet ELISA kit (R&D Systems) according to the manufacturer’s instructions. ELISA plate optical density was measured at 570nm and subtracted from readings at 450nm using a microplate reader. Cytokine levels were interpolated from a standard curve.

### SDS-PAGE and Western blot

Proteins in cell supernatants and lysates were precipitated by methanol/chloroform precipitation and resuspended in 50uL of 1X SDS-PAGE sample buffer (0.1% β-mercaptoethanol, 0.0005% bromophenol blue, 10% glycerol, 2% SDS, 63mM Tris-HCl pH 6.8). After a 10-minute incubation at 95°C, samples were resolved by SDS-PAGE, transferred onto a PVDF membrane, blocked in 5% non-fat dry milk in Tris-buffered saline with 0.1% Tween20 (TBS-T) for 2 hours, and probed with primary antibodies diluted in the same 5% milk solution at 4°C overnight. Following three 10-minute washes in TBS-T, membranes were probed with infrared dye-conjugated secondary antibodies diluted in 0.5% non-fat dry milk in TBS-T for one hour before three more 10-minute washes in TBS-T. Membranes were then imaged on a LI-COR Odyssey Infrared Imaging System and visualized using ImageJ software. Antibodies and their working dilutions were as follows: rabbit-α-human NLRP3 at 1:1000 (Cell Signaling, 13158), mouse-α-human IL-1β at 1:1000 (Cell Signaling, 12242), rabbit-α-human Caspase-1 at 1:1000 (Cell Signaling, 2225), IRDye 800CW goat-α-mouse IgG at 1:5000 (LICORbio), IRDye 800CW goat-α-rabbit IgG at 1:5000 (LICORbio).

### Statistics

All data were plotted and analyzed using GraphPad Prism 10 software and are presented as the mean ± standard error of mean (SEM). *N*-values in the figures refer to biological replicates and at least 3 replicates were conducted per condition and experiment. *P*-values were calculated using Student’s t-tests for two groups. Statistical significance for experiments with more than two groups was tested with two-way ANOVA with multiple comparison test correction as indicated.

## Results

### *TOMM40* knockdown inhibits NLRP3 activation by inhibiting NLRP3 inflammasome pathway gene expression

To determine the effect of *TOMM40* KD on NLRP3 activation, we differentiated THP-1 monocytes into macrophages and transfected them with *TOMM40*-targeting or non-targeting control (NTC) siRNAs for 48 hours (h), achieving 40% *TOMM40* KD (**Figure 1A**). We then primed them with bacterial lipopolysaccharide (LPS) to produce NLRP3 monomers and pro-IL-1β before stimulating them with oxLDL or the NLRP3 agonist, Nigericin, for 24h to induce NLRP3 assembly and subsequent IL-1β and IL-18 release. Supernatant IL-1β and IL-18 levels, measured as a readout of NLRP3 activation, were significantly reduced by *TOMM40* KD (**Figure 1B**). While a slight reduction in IL-1β release was already detected at the priming stage, this reduction became much more pronounced after stimulation with oxLDL or Nigericin. As we only saw substantial IL-18 release after stimulation with a high concentration of Nigericin (5uM) while robust IL-1β release occurred after stimulation with a lower concentration (1uM), we expected to see less IL-18 release in response to oxLDL (**Figure 1B**). However, *TOMM40* KD consistently inhibited both IL-1β and IL-18 release, supporting the conclusion that *TOMM40* KD inhibits NLRP3 activation specifically rather than through other *de novo* mechanisms of IL-1β release.

**Figure 1.**
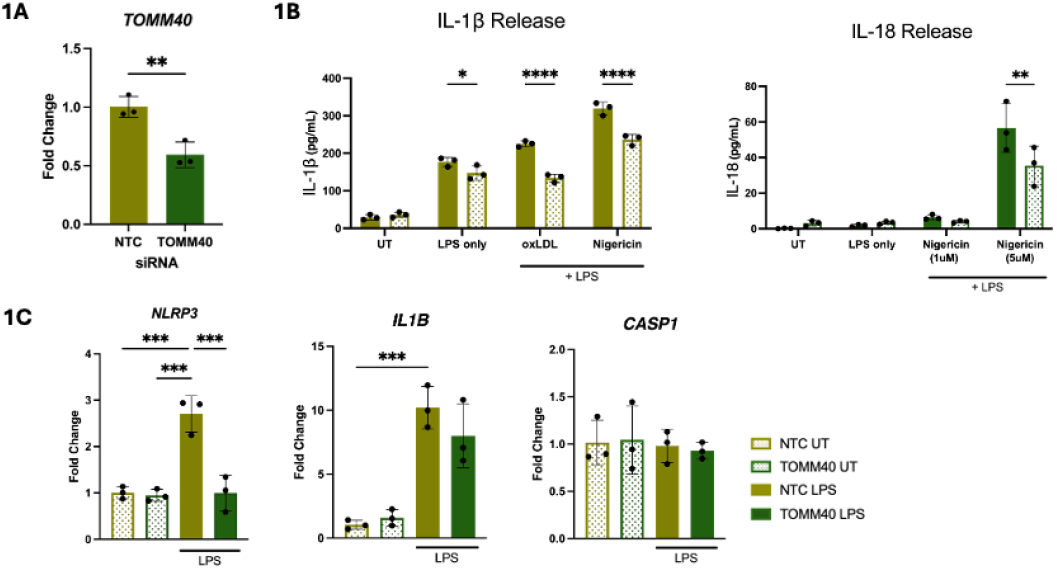
*TOMM40* KD inhibits NLRP3 activation in THP-1 monocyte-derived macrophages. **(A-C)** THP-1 monocytes were differentiated into macrophages via phorbol 12-myristate 13-acetate (PMA) treatment before transfection with a non-targeting control (NTC) siRNA or *TOMM40*-targeting siRNA for 48h. Cells were then primed with LPS for 4h and stimulated with 20ug/mL oxLDL or 1-5uM Nigericin, a NLRP3 inflammasome activator. After 24h, supernatants and pellets were collected. (**A&C**) RNA extraction and cDNA synthesis was performed on cell pellets before gene expression levels were measured by qRT-PCR. Fold changes in transcript levels are shown for *TOMM40, NLRP3, CASP1, and IL-1β* between NTC and *TOMM40* KD groups treated as indicated. (**B**) IL-1β & IL-18 levels in supernatants were quantified by ELISA. Solid bars indicate NTC while dotted bars indicate TOMM40 KD. ns= not significant, *p<0.05, **p<0.01, ***p<0.001, ****p<0.0001

To assess whether *TOMM40* KD affects NLRP3 activation at the upregulation stage or assembly stage, we next examined *NLRP3, CASP1,* and *IL1B* mRNA expression levels after LPS priming. We found that *NLRP3* and *IL1B* were upregulated in primed macrophages as expected, but *NLRP3* upregulation was abolished by *TOMM40* KD (**Figure 1E**). While there was a modest reduction in *IL1B* expression (21.8%), this was not statistically significant. There were no discernible differences in *CASP1* expression in response to priming or *TOMM40* KD as expected (**Figure 1E**). We therefore conclude that TOMM40 KD likely inhibits NLRP3 activation by a transcriptional mechanism, though it may also inhibit assembly.

### *TOMM40* KD inhibits NLRP3 activation and increases cholesterol transporter gene expression in an LXR-β-modulated manner

Given that *TOMM40* KD increases LXR-β activity in hepatocytes, we hypothesized that it does the same in macrophages, which should result in decreased pro-inflammatory signaling and increased cholesterol efflux. To this end, we first tested the effect of *TOMM40* KD on *NR1H2* (*LXRB)* and *NR1H3 (LXRA)* expression, which encode for LXR-β and LXR-α respectively, and found that *LXRB* was upregulated while there was no significant effect on *LXRA* (**Figure 2A**). To determine whether *TOMM40* KD-induced NLRP3 inhibition is LXR-dependent, we assessed the effect of treatment with GSK2033, an antagonist for both LXR-α and β, on IL-1β release. We found that pre-treatment with GSK2033 for 24h reduced IL-1β release following oxLDL or Nigericin stimulation in both *TOMM40* KD and NTC macrophages and abolished the differences in IL-1β release between these cells (**Figure 2B**). These findings indicate that in activated macrophages, LXR-α and/or LXR-β mediate IL-1β release and its inhibition by *TOMM40* KD. Since only LXR-β expression is significantly altered by *TOMM40* KD, this suggests that *TOMM40* KD-induced NLRP3 inhibition is LXR-β-modulated.

**Figure 2.**
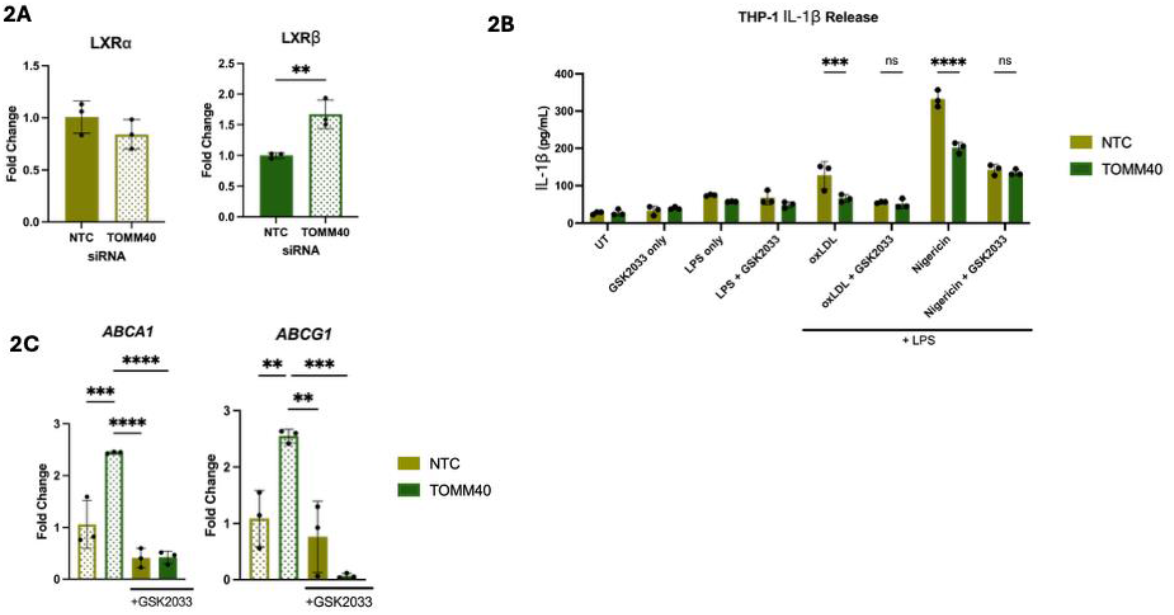
*TOMM40* KD inhibits NLRP3 activation and increases cholesterol transporter gene expression in an LXR-β-modulated manner. (A&C) THP-1 monocytes were differentiated into macrophages via phorbol 12-myristate 13-acetate (PMA) treatment before treatment with GSK2033 for 24h followed by transfection with NTC or *TOMM40*-targeting siRNAs for 48h. Cell pellets were collected and analyzed by qRT-PCR to determine fold change in transcript levels for *TOMM40, LXRA, LXRB, ABCA1, ABCG1*, and *SRB1* genes between NTC and *TOMM40* KD groups treated as indicated. (**B**) 24h before stimulation with oxLDL or Nigericin, cells were treated with 5uM of the LXR antagonist GSK2033. Supernatant IL-1β levels were measured by ELISA. ns= not significant, *p<0.05, **p<0.01, ***p<0.001, ****p<0.0001

We also examined whether the increase in *LXRB* expression resulted in greater LXR-β activity by assessing expression of the LXR target genes *ABCA1* and *ABCG1*, which encode transporters that mediate cellular cholesterol efflux, and found that both were upregulated (**Figure 2C**). We further found that pre-treatment with GSK2033 abolished *TOMM40* KD-induced upregulation of these genes (**Figure 2C**). There were no significant differences in *ABCA1* or *ABCG1* expression between untreated and GSK2033-treated, NTC-transfected macrophages. These findings suggest that *TOMM40* KD may increase macrophage cholesterol efflux via LXR-β activation.

## Discussion

We here show that KD of *TOMM40*, a gene encoding a key protein of the mitochondrial TOM complex, inhibits oxLDL-induced NLRP3 activation in macrophages, a process that is critical for atherogenesis. Further, we demonstrate that in macrophages, as previously shown in hepatocytes (31), *LXRB*, but not *LXRA*, is upregulated in macrophages by *TOMM40* KD, and that pharmacologic inhibition of LXR activity abolishes the effect of *TOMM40* KD on NLRP3 activation. Our previous studies in human hepatoma cells showed that *TOMM40* KD resulted in disruption of mitochondria-endoplasmic reticulum contact sites, with resulting accrual of reactive oxygen species (ROS) and non-enzymatically derived oxysterols that trigger LXRβ activation (31). It is therefore notable that despite its induction of potentially inflammatory ROS, *TOMM40* KD results in an overall inhibition of NLRP3 activation.

While we find that *TOMM40* KD inhibits *NLRP3* upregulation in response to LPS priming in an LXR-dependent manner, further investigation is needed to identify the molecular mechanism for this effect. Previous studies have shown that LXR has binding affinity for pro-inflammatory gene enhancers with NF-κB binding sites that are known to be epigenetically-opened post-LPS exposure, and since *NLRP3* upregulation is also NF-κB-dependent, it may be that *TOMM40* KD-induced *LXRB* upregulation prevents NF-kB binding necessary for *NLRP3* transcription^34–36^.

Further work is also needed to determine whether *TOMM40* KD also impacts NLRP3 assembly or if inhibition of NLRP3 assembly and subsequent cytokine release is solely a result of *NLRP3* transcriptional upregulation.

In addition to its anti-inflammatory effect, we find that *TOMM40* KD upregulates LXR-mediated expression of *ABCA1* and *ABCG1*, genes encoding proteins that mediate cellular cholesterol efflux and hence may also retard atherogenesis by promoting reverse cholesterol transport. To this end, future studies could measure the effect of *TOMM40* KD on macrophage cholesterol content following cholesterol loading and thus elucidate whether our documented effect on transporter expression translates to increased cholesterol efflux.

In summary the present findings present evidence supporting novel potentially anti-atherogenic effects of suppressing *TOMM40* expression in macrophages. Future studies aimed at documenting these effects *in vivo* would support the rationale for identifying therapeutic agents that target this pathway. Despite the well-established physiologically beneficial effects of LXR activity on cellular cholesterol efflux and inflammatory mechanisms, LXR agonists in clinical trials have failed to progress as increased LXR activity also induces expression of lipogenic genes which leads to hypertriglyceridemia and even steatohepatitis^38,41–43^. However, studies in mice find that these adverse effects of LXR activation are primarily due to LXR-α activity and thus there is significant interest in an LXR-β specific agonist to prevent or even treat atherosclerosis^44–46^.

## Acknowledgements

We thank Marisa Medina for her support throughout this project, Sarah King for her helpful guidance and administrative support, and Yuqing Zhang, Tommy Tran, and Shlok Rajurkar for their experimental assistance.

